# Machine learning-guided acyl-ACP reductase engineering for improved *in vivo* fatty alcohol production

**DOI:** 10.1101/2021.05.21.445192

**Authors:** Jonathan C. Greenhalgh, Sarah A. Fahlberg, Brian F. Pfleger, Philip A. Romero

## Abstract

Fatty acyl reductases (FARs) catalyze the reduction of thioesters to alcohols and are key enzymes for the microbial production of fatty alcohols. Many existing metabolic engineering strategies utilize these reductases to produce fatty alcohols from intracellular acyl-CoA pools; however, acting on acyl-ACPs from fatty acid biosynthesis has a lower energetic cost and could enable more efficient production of fatty alcohols. Here we engineer FARs to preferentially act on acyl-ACP substrates and produce fatty alcohols directly from the fatty acid biosynthesis pathway. We implemented a machine learning-driven approach to iteratively search the protein fitness landscape for enzymes that produce high titers of fatty alcohols *in vivo*. After ten design-test-learn rounds, our approach converged on engineered enzymes that produce over twofold more fatty alcohols than the starting natural sequences. We further characterized the top identified sequence and found its improved alcohol production was a result of an enhanced catalytic rate on acyl-ACP substrates, rather than enzyme expression or *K*_*M*_ effects. Finally, we analyzed the sequence-function data generated during the enzyme engineering to identify sequence and structure features that influence fatty alcohol production. We found an enzyme’s net charge near the substrate-binding site was strongly correlated with *in vivo* activity on acyl-ACP substrates. These findings suggest future rational design strategies to engineer highly active enzymes for fatty alcohol production.

## Introduction

Fatty acyl reductases (FARs) are vital for the microbial synthesis of key primary and secondary metabolites such as fatty aldehydes, waxes, alkanes, and fatty alcohols. These enzymes often interface with fatty acid anabolic/catabolic pathways and catalyze the reduction of thioester bonds found in acyl-acyl carrier proteins^1^ (acyl-ACPs) and acyl-coenzyme As (acyl-CoAs)^2^. These enzymes typically have a preference for either acyl-ACP or acyl-CoA substrates, but also display cross reactivity due to the common thioester bond in both substrates. Some FARs perform only one, two-electron, reduction step to produce aldehydes^3^, while others can perform two sequential reduction steps (totaling four electrons) to produce alcohols directly^4–6^.

The alcohol-forming FAR enzymes capable of complete reduction of thioesters to alcohols have been widely used in metabolic engineering for producing fatty alcohols^7–11^. The enzymes Maqu 2220 and MA-ACR from *Marinobacter aquaeloei* display high activity on acyl-CoA substrates and produce the corresponding fatty alcohols^2,4,11^. These enzymes can be incorporated to feed off of the reverse beta oxidation pathway to yield high levels of alcohols^8^. Another common metabolic engineering strategy involves terminating the host organism’s fatty acid elongation cycle with a thioesterase to produce a fatty acid that can then be converted to an acyl-CoA by an ATP dependent ligase, and then finally converted to an alcohol by a FAR^7,9,12^. This approach was recently applied using an engineered C8-specific thioesterase to produce octanol at a titer of 1.3 g/L^9^. While these titers are impressive, alcohol production could be more efficient with enzymes that bypass the thioesterase-ligase route, and instead directly convert acyl-ACPs to alcohols^13^.

Alcohol-forming FARs that prefer acyl-ACP substrates are less well characterized, and often display low to moderate activity relative to enzymes that prefer acyl-CoA substrates. Engineering alcohol-forming FARs such as MA-ACR to have higher activity on acyl-ACP substrates would open up new highly efficient pathways to making fatty alcohols *in vivo*. However, these enzymes are challenging to engineer using traditional protein engineering methods. MA-ACR and its close homologs lack high-resolution crystal structures needed for most computational and rational engineering approaches. Directed evolution strategies are also difficult because fatty alcohol production cannot be assayed in high throughput. Machine learning (ML)-based protein engineering has recently emerged as an efficient strategy for engineering proteins with limited structural and functional information^14–20^. Machine learning algorithms can infer the protein sequence-function mapping given a limited experimental sampling of the landscape^14^. The resulting sequence-function models can be used to computationally explore sequence space and predict optimized sequences.

In this work, we apply an ML-based protein engineering framework to engineer acyl-ACP reductases to produce fatty alcohols *in vivo*. We start by characterizing the ability of MA-ACR and related enzymes to produce fatty alcohols from intracellular acyl-ACP pools. We then design a large library of chimeric enzymes and develop an ML-based protein optimization strategy to rapidly identify highly active sequences. Our approach consists of generating diverse initial sequence sampling to get a preliminary view of the landscape, followed by iterative design-test-learn cycles to efficiently search the landscape and discover optimized sequences. After ten design-test-learn cycles, the algorithm converged on highly active acyl-ACP reductases that produce 4.9-fold more fatty alcohols than MA-ACR. We evaluated the performance of the engineered enzymes *in vitro* and found the improved alcohol titers are the result of engineered enzymes with increased catalytic efficiency. Finally, we performed a statistical analysis of the landscape and identified key sequence elements that contribute to enzyme activity. Many of these elements are located near the enzyme’s putative substrate entry channel and may be involved with modulating the preference between acyl-CoA and acyl-ACP substrates. These results open future directions to engineer enzymes for efficient microbial production of fatty alcohols.

## Results

### In vivo fatty alcohol production by natural and chimeric acyl-ACP reductases

We focused our protein engineering efforts on MA-ACR from *Marinobacter aquaeloei* because it displays high *in vivo* activity on acyl-CoA substrates^7–9^ and it was also suspected to accept acyl-ACP substrates. MA-ACR consists of two domains that sequentially reduce thioesters to alcohols (Figure 1a). The C-terminal acyl-thioester reductase (ATR) domain reduces thioesters from ACP or CoA substrates to aldehydes, and the N-terminal aldehyde reductase domain (AHR) reduces aldehydes to alcohols^4^. We also identified two related enzymes from *Marinobacter BSs20148* and *Methylibium Sp. T29* that have 60-81% sequence identity with MA-ACR (Figure 1b) and were previously shown to produce alcohols from acyl-CoAs^8,9^. Throughout the remainder of this paper, we refer to the FAR enzymes from *Marinobacter aquaeloei, Marinobacter BSs20148*, and *Methylibium Sp. T29* as MA-ACR, MB-ACR, and MT-ACR, respectively.

**Figure 1:**
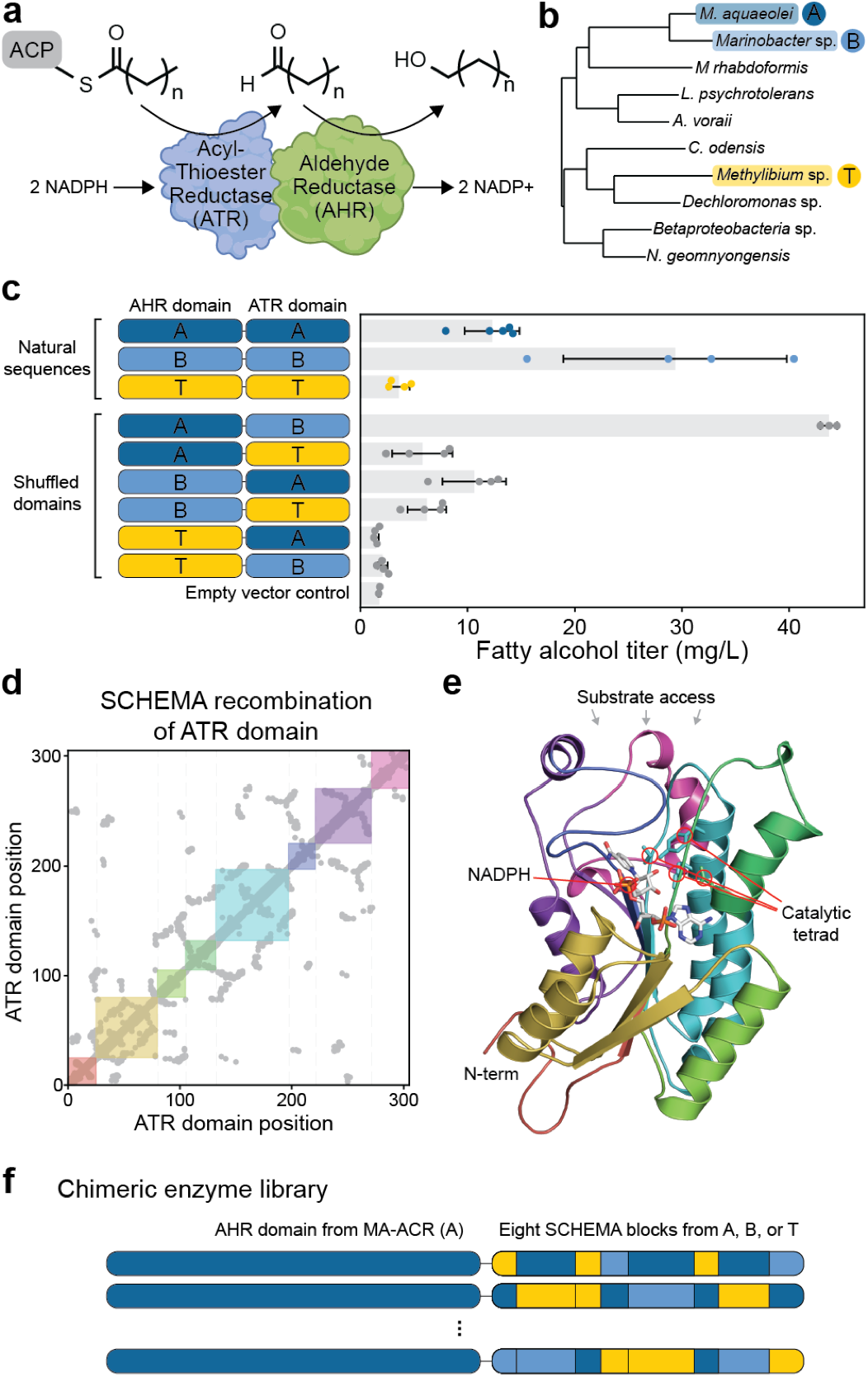
Acyl-ACP reductase activity of natural and chimeric enzymes. (a) Alcohol-forming acyl-ACP reductases consist of two domains that sequentially reduce acyl-ACP substrates to aldehydes, and then aldehydes to alcohols. (b) We focused our studies on three diverse sequences from *M. aquaeloei, Marinobacter BSs20148*, and *Methylibium Sp. T29*, which we refer to as A, B, and T, respectively. (c) Total fatty alcohol production by the three natural sequences and the six chimeric enzymes generated by shuffling their AHR and ATR domains. (d) ATR domain residue-residue contact map used for SCHEMA recombination. The colored squares depict the eight sequence blocks from the SCHEMA design that minimizes structural disruption. (e) The SCHEMA blocks mapped onto the ATR domain’s three-dimensional structure. (f) Our chimeric ATR library was fused to the AHR domain from MA-ACR.

We characterized the ability of these three natural enzymes to produce fatty alcohols from intracellular acyl-ACP pools by introducing them into *E. coli* RL08ara^21^, a strain that lacks the *fadD* gene, which encodes an acyl-CoA ligase. Deletion of *fadD* decreases the formation of acyl-CoAs and thus present the enzymes with substrates that are predominantly acyl-ACPs from fatty acid biosynthesis^10,13^. We grew each strain under aerobic conditions, extracted the fatty alcohols and measured the fatty alcohol (C6-C16) titers using gas chromatography. We found enzyme MB-ACR from *Marinobacter BSs20148* displayed more than double the total fatty alcohol titer of MA-ACR (Figure 1c). These results suggest that MB-ACR may have a preference for acyl-ACP substrates because it was previously shown to have lower activity than MA-ACR on acyl-CoA substrates^8^.

We next characterized the fatty alcohol production from chimeric enzymes generated by swapping AHR and ATR domains between the three natural sequences. Of the six possible chimeric enzymes, we found the chimera with an AHR domain from MA-ACR and the ATR domain from MB-ACR displayed the highest fatty alcohol titers (Figure 1c). This chimeric enzyme produced ∼50% more fatty alcohol than MB-ACR and roughly three-fold more fatty alcohol than MA-ACR. The ATR domain from MT-ACR also displayed increased activity (∼1.5x) when fused to the AHR domain from MA-ACR. These results suggest that MA-ACR’s AHR domain is more efficient than the AHR domains from the two other natural enzymes.

To further explore how gene shuffling can enhance fatty alcohol production, we designed a large library of ATR domains using SCHEMA^22–24^ structure-guided recombination (Figure 1d). Our design used a homology model of MA-ACR’s ATR domain to define the family’s contact map and identified seven breakpoints within the domain that balance structural disruption with library diversity (Figure S1). These seven breakpoints define eight sequence blocks that span the ATR domain’s structure (Figure 1e). Notably, the structure’s substrate access channel is composed of blocks 4, 5, 6, 7, and 8, and diversity at these positions may result in changes in the enzyme’s substrate preference. Each of the eight sequence blocks can be inherited from one of the three natural enzymes to define a combinatorial sequence space of 3^8^ sequences. However, block 6 from MA-ACR and MB-ACR happened to be perfectly conserved, and therefore the total library diversity is 2*3^7^ = 4,374 sequences. We fused our chimeric ATR domains with the highly active AHR domain from MA-ACR (Figure 1f).

For the remainder of the paper, we refer to chimeras by a block sequence (e.g. A-ABTABTAB) that specifies which of the three enzymes each sequence fragment was inherited from. Here, A, B, and T correspond to MA-ACR, MB-ACR, and MT-ACR, respectively; the first position specifies the AHR domain and the remaining positions specify the ATR domain’s eight SCHEMA blocks. We also refer to the three sequences that have all eight ATR blocks from a single natural enzyme as “parental” enzymes. Here “parent A” has the block sequence A-AAAAAAAA, “parent B” is A-BBBBBBBB, and “parent T” is A-TTTTTTTT.

### Increasing fatty alcohol production with ML-driven enzyme engineering

We aimed to identify the most highly active enzymes from our chimeric ATR domain library. However, the chimera space consists of thousands of unique sequences and is much too large to fully characterize using our low throughput gas chromatography assay. Instead, we developed an ML-based sequence optimization method to rapidly identify highly active sequences with minimal experimentation (Figure 2a). Our approach consists of a generating diverse initial sequence sampling to get a preliminary view of the landscape, followed by iterative design-test-learn cycles to efficiently search the landscape and discover optimized sequences.

**Figure 2:**
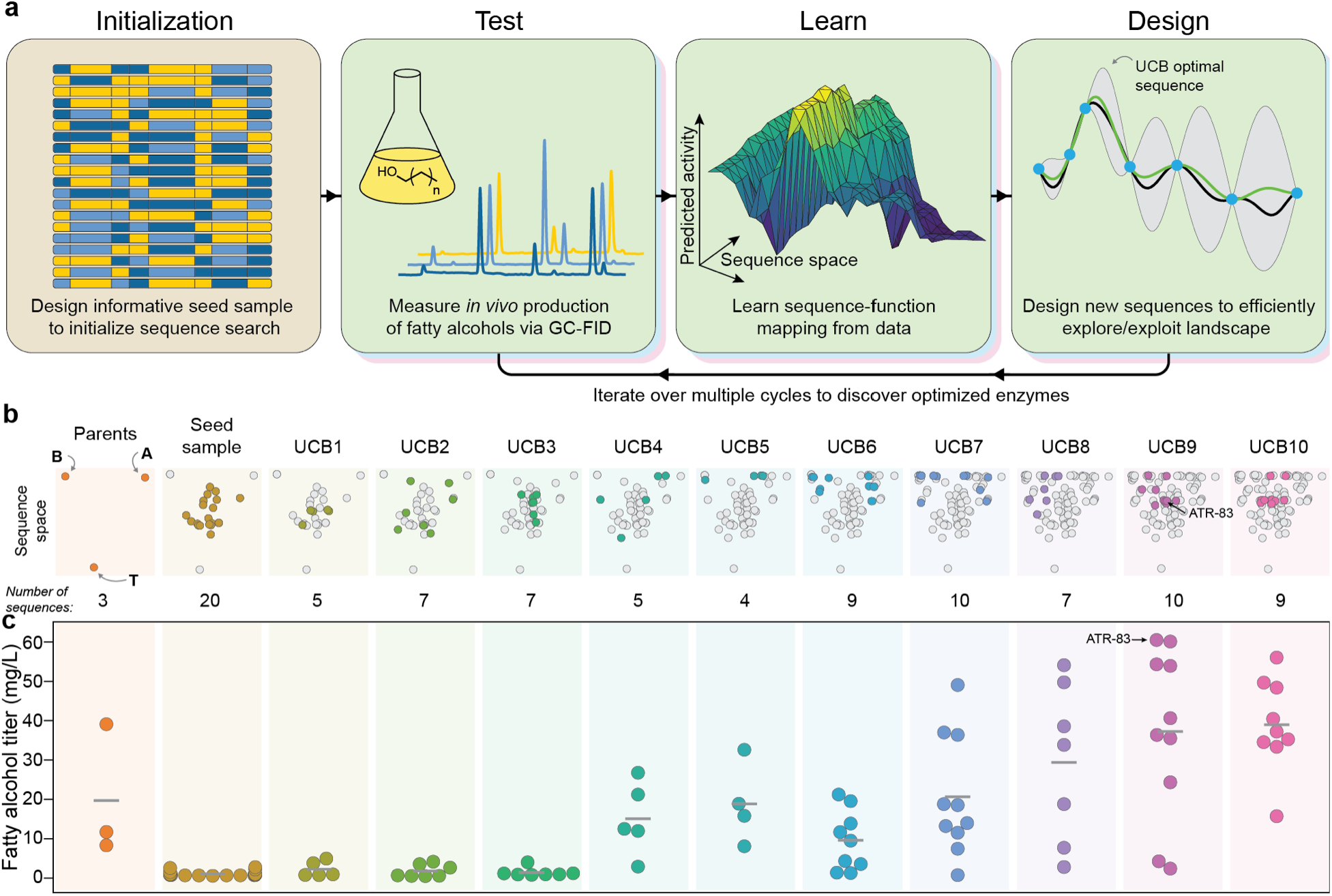
ML-accelerated protein sequence optimization. (a) An overview of our sequence space search strategy. We first initialize the search by designing a diverse set of sequences that broadly sample then landscape. We then iterate through multiple design-test-learn cycles to efficiently understand and optimize *in vivo* fatty alcohol production. (b) Sequence space visualization over ten rounds of UCB optimization. The three parent enzymes are found at the vertices of this chimeric sequence space and all chimeras fall within the parents’ envelope. The UCB optimization started by broadly sampling the landscape, but quickly converged on highly active regions. (c) The *in vivo* fatty alcohol titers over the course of the sequence optimization. Each point depicts an individual sequence’s fatty alcohol production and the horizontal grey bars represent the average titer during that round of sequence optimization.

We generated a diverse initial sampling of sequence space by identifying the 20 sequences that are maximally representative of the full chimera space consisting of 4,374 sequences. We then constructed these sequences and experimentally measured their fatty alcohol titers (Figure S2). Seventeen of these 20 sequences displayed no measurable alcohol production and the remaining three produced low titers that were below the least productive parent (T). The fatty alcohol titer data from these 20 initial sequences was used to train a Gaussian process (GP) sequence-function model that can make predictions across the entire chimera space (Figure S3). Importantly, GPs also provide estimates of the model’s uncertainty (confidence intervals) that can be used to gauge the reliability of predictions and highlight gaps in its understanding of the landscape^14,15^.

With the initialized GP sequence-function model, we then iterated through multiple design-test-learn cycles with the goal of identifying the optimal sequence with minimal experimental samples. The sequences for the next round of experimentation were designed using an upper-confidence bound (UCB) criterion that simultaneously explores uncertain regions of the landscape and samples sequences that are predicted to be optimized. UCB optimization provides strong theoretical guarantees for efficiently balancing exploration and exploitation^25,26^, and should rapidly converge on the optimal sequences. During each iteration, we designed 5-12 sequences using the UCB criterion, assembled the corresponding genes, transformed them into *E. coli*, and measured each strain’s fatty alcohol titer using gas chromatography. The new data was then used to update the sequence-function model and the process was repeated. We performed a total of ten rounds of UCB sequence optimization and saw gradual improvements in fatty alcohol titers (Figure 2). The details of each round of UCB optimization can be found in Supplemental Table 4.

The UCB sequence optimization converged on multiple highly active acyl-ACP reductases. The enzyme with the highest titer had a block sequence of A-ATBBAAAB and we refer to this top sequence as ATR-83. Additional *in vivo* characterization showed that ATR-83 produces a total titer of 54 ± 11 mg/L fatty alcohols (Supplementary Table 5), which is nearly five-fold greater than the titer of MA-ACR and about two-fold greater than the best natural sequence (MB-ACR). The alcohols produced by ATR-83 and the other top chimeras consisted of primarily hexadecanol (C16) and some tetradecanol (C14). This product distribution is expected since long chain acyl-ACPs are the primary precursors for the lipids that make up the cell membranes in *E. coli*^27,28^.

### Improved fatty alcohol production occurs via an enhanced catalytic rate on acyl-ACP substrates

Our engineered acyl-ACP reductase chimeras produce several fold more fatty alcohols than the initial natural sequences. Increased flux through the metabolic pathway can be the result of improved protein stability and/or expression, enzyme kinetic properties, or possibly interactions with other components of the pathway. We performed further biochemical analysis of the engineered enzymes to better understand how they increase alcohol production.

We first measured the level of enzyme expression in the production strain (Figure S4, Figure 3a). We found all sequences were expressed at high levels and there were no statistically significant differences between the natural and engineered sequences. Next, we purified the enzymes and measured their kinetic properties on palmitoyl-ACP (Figure 3bc). The four enzymes tested displayed similar *K*_*M*_ values for palmitoyl-ACP, but ATR-83 had a substantially larger turnover number. ATR-83’s increase in *k*_*cat*_ matches its improvements in fatty alcohol titer. Taken together with the enzyme expression data, this suggests that the engineered enzymes are increasing alcohol production by an enhanced catalytic rate.

**Figure 3:**
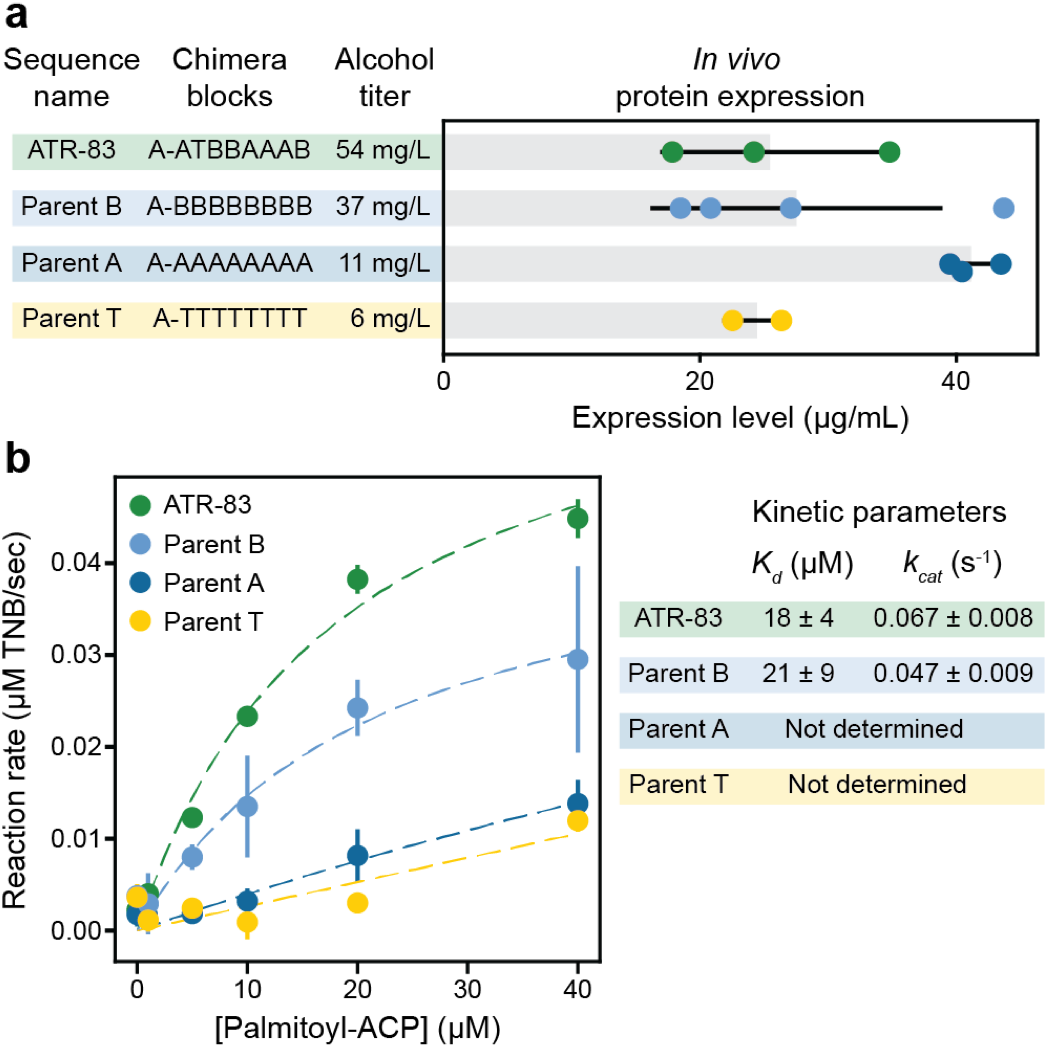
Expression levels and kinetic activity of selected ATRs. (a) We measured the expression levels of the ATR-83 chimera and the three parental enzymes. These four enzymes displayed no significant differences in expression despite the large differences in their alcohol titer. (b) We characterized the kinetics of selected enzymes on palmitoyl-ACP. ATR-83 displayed a higher turnover number (*k*_*cat*_) relative to the three parental enzymes. The kinetic parameters for parents A and T could not be precisely determined due to their low overall activity and the resulting poor fit to the Michaelis-Menten model.

We also analyzed the enzymes’ activity on CoA substrates and found that ATR-83 has a lower activity than the parents on palmitoyl-CoA (Supplemental Figure S5). This suggests that ATR-83 may not be a faster enzyme overall, but instead displays an altered preference for ACP over CoA. This altered preference could be the result of changes in the protein surface that interacts with ACP substrate.

### Statistical analysis of the enzyme landscape reveals features that influence fatty alcohol production

Over the course of our UCB sequence optimization, we collected 96 data points mapping chimeric sequences to fatty alcohol titers. This sequence-function data can serve as a rich resource for understanding how protein sequence and structure impact *in vivo* enzyme activity. We trained a GP regression model to predict fatty alcohol titers from sequence. This model displayed excellent predictive ability in a cross-validation test (Figure S6).

We used this predictive model to assess how each chimera sequence block contributes to overall enzyme activity (Figure 4a). We see that most block positions influence activity and display a broad range of effects. The three sequence blocks with the largest positive contribution were block 7 from MA-ACR, block 3 from MB-ACR, and block 2 from MT-ACR. Substitution to any one of these blocks tends to increase alcohol titers by over 70%. Block 8 from MB-ACR also strongly tends to increase the titers. The sequence blocks with the most negative contribution were blocks 3 and 7 from MT-ACR. Overall, most blocks from MT-ACR were deleterious for alcohol production.

**Figure 4:**
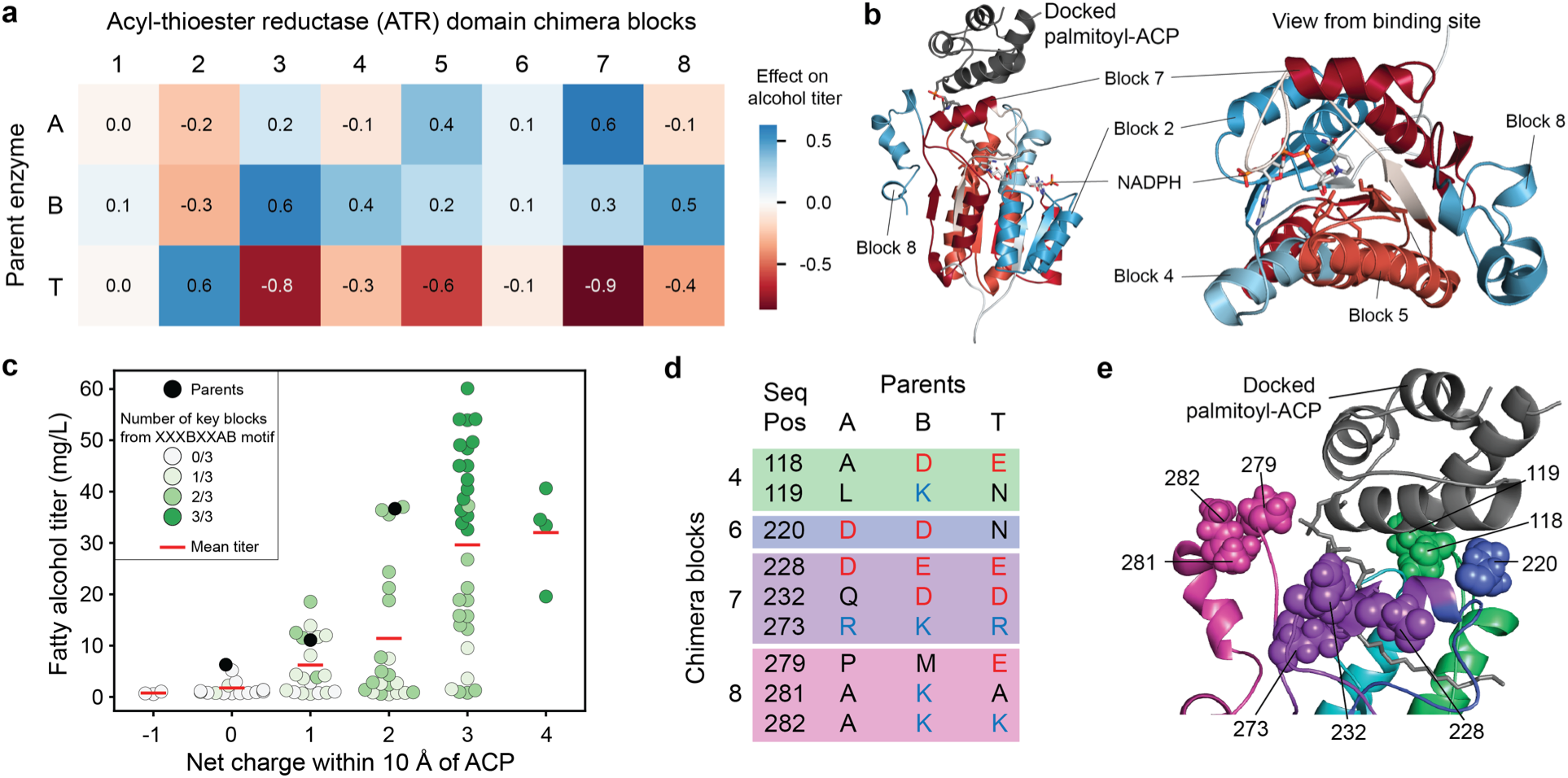
Statistical analysis of the fatty alcohol production landscape. (a) Contributions of each sequence block to the alcohol production of chimeras. (b) Mapping of the contributions to the structural model of MA-ACR. Blocks with strong effects (either positive or negative) line up with key structural features such as the NADPH binding domain, the active site and the ACP binding site. (c) Correlation between net charge near the binding site and fatty alcohol titer of chimeras. Chimeras with higher net charges tend to produce higher titers. Additionally, three key blocks were found to correlate with both activity and with charge. Combining all three of these blocks results in highly active enzymes. (d) Positions in the parent sequence alignment that contain non-conserved charged residues within 10 Å of the putative ACP binding site. (e) Locations of key charged residues in the structural model of MA-ACR docked with palmitoyl-ACP. Optimal combinations of blocks could produce more favorable interactions with the ACP.

We mapped the block effects onto MA-ACR’s homology model to relate their contributions to structure and mechanism (Figure 4b). Block 2 likely forms extensive interactions with the enzyme’s NADPH cofactor, and MT-ACR is the best parent at this position. While there are many amino acid differences in this block, it’s notable that MT-ACR has a different NADPH binding motif than the other two parents (GGSSGIG vs GATSGIG). MT-ACR’s motif may provide more efficient NADPH utilization *in vivo*. Blocks 4-8 make up the binding pocket for the acyl-thioesters. Block 5 contains three of the catalytic residues (a Y, S and K), and block 6, whose sequence is highly conserved, appears to be involved in NADPH binding. Blocks 7 and 8 appear to contain surface residues; positively charged residues in these blocks are likely involved in docking the negatively charged acyl-ACP^29^.

We hypothesized the net charge of the enzyme’s substrate binding pocket may influence activity because the ACP substrate contains many negatively charged residues. To examine the enzyme’s charge distribution near the substrate binding site, we computationally docked ACP (from PDB entry 6DFL) to our homology model of MA-ACR using RosettaDock^30^. We then identified all interface positions within a 10 Å radius of the docked ACP and calculated the net charge of each chimera’s interface residues. We found the net charge of an enzyme’s substrate binding interface was positively correlated with the total fatty alcohol titer (Figure 4c). A chimera’s substrate interface charge is dictated by seven sequence positions that are near the ACP substrate and that also display variation in charged residues among the parents (Figure 4de). The charges at these sequence positions can largely explain the preferred blocks from Figure 4a.

## Discussion

Engineering fatty acyl reductases (FARs) to have improved activity on acyl-ACP substrates could open new routes to *in vivo* production of fatty alcohols, and other valuable bioproducts such as waxes and alkanes. In this work, we engineered enzymes with improved activity on acyl-ACP substrates. Our approach leveraged gene shuffling to broadly sample sequence space and ML-driven protein engineering to rapidly and efficiently identify optimized sequences. Our top identified enzyme, ATR-83, displayed two-fold higher *in vivo* fatty alcohol titers than the best natural sequence MB-ACR and nearly five-fold higher titers than MA-ACR. These increases in fatty alcohol titer are a result of ATR-83’s enhanced turnover number on ACP substrates. The chimeric enzymes discovered in this work have potential to improve the efficiency of alcohol production from acyl-ACPs *in vivo*.

Shuffling the AHR and ATR domains between the three natural sequences generated chimeric enzymes that produce a broad range of fatty alcohol titers. From these results, it appears the ATR domain from MB-ACR has the highest activity on ACP substrates and the AHR domain from MA-ACR has the highest activity on the intermediate aldehyde substrate. Rather than directly affecting the catalytic rate, it’s also possible that these domains could be enhancing activity through inter-domain interactions, especially since MA-ACR has been shown to be tetrameric^4^.

Machine learning is rapidly advancing the fields of directed evolution and protein engineering^15–17,31^. Though some ML-based strategies (especially those involving deep learning or neural nets) require massive amounts of training data, active-learning approaches (such as UCB optimization) can be used to simultaneously explore the sequence-function landscape and identify improved sequences from relatively few data points. The reduced need for data enables protein engineering workflows that don’t depend on high-throughput techniques, and thus overcomes major limitations of directed evolution approaches. Our design-test-learn cycle closely resembles the UCB optimization process previously used to engineer thermostable chimeric cytochrome P450s^14^. However, a key difference in this work was the introduction of an active/inactive binary classifier to filter out potential inactive sequences that provide little information regarding enzyme activity. Incorporating this classifier led to improved predictions by the GP regression model, especially in early UCB rounds when the number of active sequences was small (only 12 sequences were active from the first three rounds).

In the early rounds of our UCB sequence optimization, we found it was helpful to restrict the number of block exchanges from the parent sequences in order to bias the search towards functional sequences. Sampling further away almost always resulted in non-functional sequences that provided little information about the fatty alcohol production landscape. We learned this trick during the course of the sequence optimization, which certainly limited the efficiency of our method. Future improvements to UCB algorithm could include an informative prior for the active/inactive binary classifier that encodes a preference to sample near the parent sequences when limited functional data is available.

ATR-83 produced 50% more fatty alcohols than parent B (A-BBBBBBBB) and 450% more than MA-ACR. It is difficult to interpret these *in vivo* results because intracellular acyl-ACP pools exist as a broad mixture from C4-C18, and each enzyme may have its own substrate preferences. We performed further kinetic characterization on the enzymes and found ATR-83’s increased *in vivo* alcohol production is the result of enhanced turnover number (*k*_*cat*_) on ACP substrates, rather than enzyme expression or *K*_*M*_ effects. Interestingly, ATR-83 displays lower activity on acyl-CoA substrates than parent B and MA-ACR. Since both acyl-ACP and acyl-CoA substrates have the same thioester bond that is being reduced, one might expect substrate specificity to manifest as differences in *K*_*m*_ between the enzymes. However, we observed enzymes’ *k*_*cat*_ to be the major determinant of substrate specificity. One possible explanation for the observed behavior could be that ACP is interacting with the enzyme surface to allosterically enhance the catalytic rate. Similar allosteric modulation by ACPs has been observed in the LovD enzyme^32^.

We found a positive correlation between an enzyme’s net charge near the putative substrate binding site and its activity on acyl-ACPs *in vivo*. This relationship may be expected because positive charges on the enzyme surface could enhance electrostatic interactions with the negatively charged ACP substrate. The chimeric enzymes’ substrate interface charge is largely dictated by blocks 4, 7, and 8. Sequences with B at block 4, A at block 7, and B at block 8 (i.e. XXXBXXAB) can increase the net interface charge of a chimera by up to +4. The average alcohol titer of chimeras containing these three blocks is 42 mg/L, compared to an average of 8 mg/L for sequences without that combination. These results suggest future enzyme engineering directions to supercharge the substrate interface with positively charged residues to further enhance electrostatic interactions with ACP. A similar approach has been applied to acyl-ACP thioesterases, leading to improved enzyme activity^29^.

While we demonstrated that our engineered enzymes have improved activity on palmitoyl-ACP both *in vivo* and *in vitro*, the activity of the enzymes on shorter and medium chain substrates is less clear. Production of medium chain fatty alcohols, such as octanol, remain a prime target for metabolic engineering, since medium chain fatty alcohols are more valuable than long chain fatty alcohols^33^. In order to explore the *in vivo* activity of these engineered enzymes on shorter chain acyl-ACPs, new methods would be needed to alter the acyl-ACP distribution in the cells without significantly disrupting pathways involving production of lipids for the cell membranes. Alternatively, pathways that utilize acyl-CoA pools show promise for making medium length alcohols selectively^7,9^. While our active-learning strategy focused on acyl-ACP activity, it could also be used to enhance activity on medium chain acyl-CoAs.

Our ability to engineer microbes to produce high-value chemicals is often limited by the availability of enzymes to catalyze key chemical reactions. We have presented an enzyme engineering framework that leverages ML-based sequence-function models with iterative experimentation to rapidly identify improved enzymes. This approach can be generally applied to enzymes that lack a high-throughput functional assay or structural information, and therefore are challenging to engineer using traditional directed evolution and rational methods. Future advances in enzyme engineering will open entirely new routes to produce valuable chemicals from low-cost and renewable feedstocks.

## Materials and Methods

### Chemicals, Reagents, and Media

*E. coli* RL08ara^21^ and CM24^8^ assay media used for this study are the same composition as Miller LB, except with 10 g/L peptone instead of 10 g/L tryptone. CM24 media was supplemented with 1% w/v glucose, and sterile filtered using a 2 µM filter. *E. coli* RL08ara assay medium was sterilized by autoclaving. Both media were adjusted to a pH of 7.0 prior to sterilization.

Individual fatty alcohol standards were prepared at a concentration of 100 mg/mL by dissolving alcohols ranging from C3 to C17 in 200 proof ethanol. Then, alcohols were mixed to make 10 mg/mL standards of even-chain alcohols (C4, C6, C8, C10, C12, C14 and C16) and odd-chain alcohols (C3, C5, C7, C9, C11, C13, C15, C17).

### Measuring in vivo fatty alcohol titers

We measured *in vivo* alcohol titers produced by each enzyme variant using gas chromatography (GC). Overnights started in LB + Kanamycin from individual colonies from the transformation were diluted into a 50 mL culture of *E. coli* RL08ara Assay Medium + Kanamycin in a 250 mL baffled shake flask such that the final OD was about 0.01. The media had a 20% (10 mL) dodecane overlay, and we supplemented the media with 1 mL of 50% v/v glycerol. The cultures grew at 37 °C for 45 minutes at 250 rpm, and then we induced protein expression by adding 500 µL of 100 mM IPTG (final concentration 100 µM IPTG). As a control, each batch also included blank cultures that were prepared by mixing media, dodecane, glycerol and antibiotic in the same amounts as the expression cultures, but without any cells added. The expression cultures incubated for 18 hours at 30 °C after induction.

Afterwards, we cooled the expression cultures on ice to prevent evaporation. Then, we added 150 µL of 10 mg/mL odd-chain internal standard mixture to each culture flask and mixed them vigorously to make an emulsion. Immediately after mixing, we transferred 5 mL of the emulsion to a glass centrifuge tube pre-loaded with 1 mL of n-hexanes. We vortexed the tubes for 20 s, shook for 20 s, and vortexed for another 20 seconds. Then, we centrifuged the samples for about 10 minutes until the organic layer and aqueous layers separated and extracted about 900 µL of the organic layer to load into a GC vial for analysis on GC-FID.

We analyzed all GC samples using a Shimadzu Model 2010 GC-FID system with an AOC-20i autosampler and a 60 m 0.53 mm ID Stabilwax column (Restek 10658). The oven temperature program used to analyze samples from RL08ara and CM24 samples was based on Mehrer et al.^8^ and is as follows: 45 °C hold for 10 minutes, ramp to 250 °C at 12 °C/minute, hold at 250 °C for 10 minutes. In some individual experiments we shortened the hold time. Each run included standards of the odd-chain internal standard mixture and even-chain standard mixture to control for any changes in the retention times of the analytes. We estimated the concentrations of even-chain fatty alcohols by averaging the areas (A_i-1_ and A_i+1_) and concentrations (C_i-1_ and C_i+1_) of the odd-chain internal standards that bracketed the particular even-chain analyte. We used the resulting response factor to convert the area of the even-chain species (A_i_) to the original media concentration (C_i_) per the following equation:

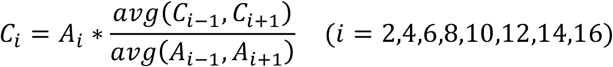

### Aerobic alcohol production in BL21(DE3)

We cloned the initial seed sample ACR chimeras into the pET28 backbone and transformed into BL21(DE3). Overnights were started in LB + Kanamycin from individual colonies from the transformation. We diluted the cultures 100-fold into 5 mL cultures of LB + Kanamycin in culture tubes. We grew the cultures for 2.5-3 hours, measured the ODs, and then induced with 5 µL of 100 mM IPTG and incubated for 24 hours at 20 °C with shaking at 250 rpm.

Following protein expression, we incubated the cultures ice for 1.5-2.5 hours. Nonanol (C9) and heptadecanol (C17) were used as internal standards; a solution that was 5 µM nonanol and 5 µM heptadecanol in hexanes was prepared and added (1 mL) to each 5 mL expression culture. We then vortexed and spun down the sample in a centrifuge (1000x G for 10 minutes) to separate the phases. 900 µL of the organic layer was extracted for analysis on GC-FID. Titers of fatty alcohols were determined using an external standard curve with standards of each of the even chain fatty alcohols in hexanes and dividing by the extraction ratio (5) to convert from the concentration in the organic phase to the original concentration in the media.

### Anaerobic alcohol production in CM24

ACR chimeras were cloned into the pBTRCK plasmid backbone and transformed into CM24 along with seFadBA (g130, pACYC-seFadBA) and tdTER (g131, pTRC99A-tdTER-fdh)^8^. We started overnights from individual colonies in LB + Kanamycin + Carbenicillin + Chloramphenicol. The following day, 600 µL of overnight cultures were diluted in 30 mL of CM24 Assay Medium + Kanamycin + Carbenicillin + Chloramphenicol with a 20 % (6 mL) dodecane overlay in a 50 mL serum vial, which was sealed. We grew the cultures for 2 hours at 30 °C, and then induced by injecting 300 µL of 100 mM IPTG (for a final IPTG concentration of approximately 100 µM) through the septum with a needle. Cultures were then incubated at 30°C for 48 hours.

Following expression, we cooled the cultures on ice and added 180 µL of an internal standard mixture (the same fatty alcohol mixture used for quantitation of alcohols in RL08ara). We mixed the samples thoroughly and extracted 5 mL of the emulsion with 1 mL of hexane per the same protocol as RL08ara above.

### Structural modeling and SCHEMA library design

We utilized the MODELLER^34^ homology modeling software to build 100 models of each of the acyl-thioester reductase domains of MA-ACR, MB-ACR, and MT-ACR using the following PDB entries as templates: 3M1A-A, 3RKR-A, 3RIH-A, 3AFM-B, 3AFN-B, and 4BMV-A. We built a contact map by determining which pairwise amino acid contacts (defined as two amino acids within a 4.5 Å radius based on any atoms in the amino acids) were present in each model, and weighted each contact by the percentage of models in which the contact was present.

We determined the crossover between the aldehyde-reductase domain and the acyl-thioester reductase (ATR) domain by aligning the sequences of MA-ACR, MB-ACR, and MT-ACR and selecting a crossover point at the conserved LDPDL, approximately 350-360 residues from the N-termini. Then, we used SCHEMA-RASPP to determine 7 additional crossover locations within the ATR domain that were compatible with Golden-Gate assembly.

### Gene assembly and strain construction

All ATR enzymes tested were cloned into the pBTRCK plasmid backbone and transformed into *E. coli* RL08ara^21^. We obtained the three natural parent sequences from prior studies^8,9^. We amplified the AHR and ATR domains of each of the natural sequences, as well as the plasmid backbone, by PCR using primers that contained Golden Gate overhangs. We used Phusion Hot Start Flex 2X Master-Mix (NEB) for all PCR reactions. Then, we used Golden Gate assembly to combine the pieces and synthesize the domain shuffled variants. Golden Gate reactions were carried out either using commercial Golden-Gate assembly mix (NEB), or an in-house mixture of the components from NEB (T4 DNA ligase buffer, BsaI HF v2 and T4 DNA ligase).

We designed plasmids containing each of the 24 blocks determined by RASPP such that each block was flanked by BsaI restriction sites. The plasmids were synthesized by TWIST Biosciences. The blocks (including the BsaI site) were amplified by PCR and cloned into a backbone vector harboring the AHR domain of MA-ACR by Golden Gate assembly. For sequences that we studied *in vitro*, we amplified the whole FAR sequence and used Golden Gate assembly to add the insert into a pET 28 backbone.

### Optimization of CM24 and BL21 UCB for UCB 1

We trained a Gaussian Process regression model on the activities from the BL21 and CM24 datasets. The trained model was then used to calculate the upper confidence bounds (UCBs) for each activity, and an algorithm was used to select sequences that maximized both UCBs. We added two UCBs together and chose the sequence with the highest summed UCB for the new set. Then, we retrained the GP model with the new sequence using the predicted mean in place of an actual experimental activity and repeated the selection process until ten new sequences were selected. Then, when five of the ten sequences were successfully assembled, we evaluated the new sequences in CM24 and RL08ara (along with all twenty sequences in the original training set being evaluated in RL08ara).

### Sequence-function machine learning and UCB optimization

To map the sequences to their function, we used a combination of Gaussian Naïve Bayes (GNB) classification and GP regression. Naïve Bayes classification aided in identifying regions of the sequence landscape that were likely to be active and allowed us to focus our predictive efforts on those regions. Then, GP regression was used to make specific predictions about the sequence-function landscape and calculate the UCB.

We used the Naïve Bayes classifier from scikit-learn to classify untested chimeras as active or inactive. We used a threshold to convert titers from the experimental data into a binary input, and then trained the classifier using those as the labels. Then, we trained a GP regression model with a linear kernel function^14^ on the titers of the sequences that were above the titer threshold. To tune the GP model and prevent overfitting, we used leave-one-out cross-validation to scan over a range of possible lambda values. We selected values that maximized the correlation coefficient and minimized the squared error; when those two objectives could not be realized simultaneously, we chose lambda values by inspection that balanced them. We then used the chosen lambda value to fit the GP model and predict the activities of all untested sequences that the GNB classifier labeled as active. We utilized batch mode UCB optimization^35^ to select 10-12 sequences to build for the next round.

### Measuring in vivo enzyme expression levels using SDS-PAGE

To verify that increases in fatty alcohol titers were due to enzyme activity, we performed additional characterization of the protein expression levels for the parents and select chimeras. To estimate the expression level of the ATR enzyme, we performed additional replicates using the same expression conditions as were used during UCB optimization. Then, after extracting the fatty alcohols, we suspended the remaining 5 mL pellet in 1 mL of media. We normalized the ODs of the suspensions to an OD of 10 and pelleted and froze 500 µL of the OD 10 culture. We later thawed the frozen pellets and lysed them using 250 µL lysis buffer (3872 µL 100 mM Tris pH 7.4, 120 µL Bugbuster, 4 µL lysozyme and 4 µL DNAse I).

We prepared a standard curve using dilutions of purified MA-ACR. We added 3 µL of each MA-ACR dilution to 12 µL of SDS master mix (which consisted of 5 parts 2X SDS mix and 1 part 1 M DTT). and mixed them in a 1:1 ratio (volume:volume) with empty vector lysate. The other lysates were mixed with 2X SDS buffer and 3 µL 100 mM Tris pH 7.4 (to ensure equal volumes of lysate between the standards and the samples). We heat denatured the lysates (at 85 °C for 2-5 minutes) and analyzed them by SDS-PAGE.

We used FIJI, an image analysis software^36^, to estimate the intensities of the ATR band in the MA-ACR standards and generate a standard curve. We made new standard curves for each replicate to reduce gel to gel variability, and only compared samples to standards on the same page gel. Expression levels are reported as µg/mL of ATR (at an OD of 20).

### Biosynthesis of fatty acyl-ACP substrates

We synthesized the acyl-ACP substrates by functionalizing purified *E. coli* ACP with a 4’-phosphopantetheine arm by the acyl-ACP synthetase from *Vibrio harveyi*^37^, and then attaching the acyl-chain to the thiol end of the arm using a phosphopantetheinyl transferase (SfP) from *Bacillus subtilis*.

### Expression of V. harveyi AasS, B. subtilis SfP and E. coli ACP

The enzymes needed to functionalize palmitoyl-ACP were expressed using the method in Hernández-Lozada et al. with some minor modifications^38^. The cells were grown for 2 hours at 37 °C (200 rpm) and then induced with 1 mM IPTG (final concentration) without cooling the cultures as was done in Hernández-Lozada et al. AasS and SfP were expressed overnight at 18 °C overnight, and ACP was expressed at 20 °C overnight and harvested by centrifugation. We also purified the proteins using the method from Hernández-Lozada et al., however rather than using dialysis, we used Amicon filter columns to carry out buffer exchange. The final concentrations of the proteins were determined using Bradford assays.

### Functionalization of E. Coli ACP

To cleave the His-tag from the Apo ACP, we added 700 uL of 2.1 mg/mL TEV protease to the 4 mL ACP solution. The reaction incubated overnight at 20 °C shaking at a speed of 250 rpm. At the conclusion of the digestion, we stored the mixture in 50% glycerol at -80 °C. Later, to purify the cleaved Apo ACP, we thawed the digestion and ran it over parallel gravity columns packed with Nickel Sepharose Fast Flow resin. We pooled the flow-through and buffer exchanged with 50 mM Na_2_HPO_4_ pH 8 + 10% glycerol using an Amicon filter unit (MWCO 3000 kDa). The concentration of the cleaved Apo-ACP was determined by a Bradford assay.

The conditions for the reactions to generate Holo-ACP were: 500 µM Apo ACP, 5 µM SfP, 5 mM Coenzyme A, and 10 mM MgCl_2_ in 100 mM Na_2_HPO_4_ pH 8. The reactions took place in 500 uL aliquots in 1.5 mL Eppendorf tubes and shaken in a beaker at 37 °C for 1 hour.

We dissolved sodium palmitate in water heated to 65 °C to a concentration of 100 mM. After the holo-ACP reactions were finished, we added palmitate, ATP, and AasS to the reaction mixture, (along with enough buffer to double the reaction volume), to give final concentrations of 5 mM palmitate, 5 µM AasS and 10 mM ATP. The reactions incubated overnight at 37 °C. Then, we pooled the reactions, purified the palmitoyl-ACP by running the mixture through a gravity column packed with Nickel Sepharose Fast Flow Resin. We buffer exchanged the purified palmitoyl-ACP into 100 mM Na_2_HPO_4_ + 10% glycerol pH 8.

### Purification of ATRs

We expressed parental ATRs (A-AAAAAAAA, A-BBBBBBBB, and A-TTTTTTTT) and purified them per the same method as *E. Coli* ACP, except for the buffer exchange step. We buffer exchanged them into 20 mM Tris, 50 mM NaCl pH 7 using an Amicon filter unit (30,000 kDa MWCO). Then, we added glycerol to the proteins (about 15 % v/v for parents 1-3). We expressed ATR-83 at 30 °C rather than 20 °C but purified it in the same manner, though we added more glycerol to the purified ATR-83 (final concentration ∼50 % v/v glycerol). We determined the concentration of the enzymes by Bradford assays.

### In vitro enzyme kinetics on palmitoyl-ACP and palmitoyl-CoA

We determined the activity of the above ATRs in a 96 well plate based assay using 5’5 Dithiobis(2-nitrobenzoic acid) or DTNB to monitor the progress of the conversion of palmitoyl-ACP to hexadecanol and free holo-ACP. We tested palmitoyl-ACP concentrations up to 40 µM (as this concentration should be within the physiological range within cells)^39^. Reactions contained 1 µM of the respective ATR and 200 µM NADPH in 20 mM Tris + 50 mM NaCl pH 7 and the total reaction volume was 100 µL. The concentration of DTNB was 250-252 µM (the difference is due to slightly different preparations of a NADPH/DTNB master mixes on different dates).

To gauge activity of the ATRs on CoAs *in vitro*, we carried out reactions using palmitoyl-CoA as a substrate. The *in vitro* assay used to determine CoA activity was identical to that used for ACP activity above.

### Computational docking and analysis of interfacial charge

We used the RosettaDock^30,40^ application to perform local docking simulations to dock a structure of palmitoyl-ACP (from PDB entry 6DFL) to MA-ACR. We did not include the acyl-chain in the docking simulations. We ran 1000 docking simulations and selected a model based on minimizing the total energy and the interface score. Then, using PyMOL, we determined which residues in the model of MA-ACR were within a 10 Å radius of the ACP molecule. The number of charged residues within that radius was then determined, and the net interface charge was defined as the number of positive residues minus the number of negative residues.

## Acknowledgements

This work was funded by the National Institutes of Health (R35GM119854) and the National Science Foundation (CBET-1703504). J.C.G. is the recipient of a National Institutes of Health Biotechnology Training Program Fellowship (NGIMS T32GM008349). The authors would also like to acknowledge Dr. Néstor Hernández Lozada, Dr. Chris Mehrer, and Dr. Mark Politz for helpful discussions regarding assay development, and Bennett Bremer for helpful discussions regarding coding and algorithms.

## Author contributions

J.C.G., B.F.P, and P.A.R conceived the project. J.C.G. performed the experiments with assistance from S.A.F.. J.C.G. analyzed the data. J.C.G. and P.A.R. wrote the manuscript with feedback from B.F.P and S.A.F.

