## Supplemental information for "Machine learning-guided acyl-ACP reductase engineering for improved *in vivo* fatty alcohol production"

**Supporting Information**

Table 1: Strain list

| Strain | Description | Source/Reference |
|---|---|---|
| *Escherichia coli* DH5α | n/a | Lucigen |
| *Escherichia coli* BL21 DE3 | n/a | Lucigen |
| *Escherichia coli* 10 G Supreme | n/a | Lucigen |
| *E. coli* RL08ara | *E. coli* K-12 MG1655 ΔfadD ΔaraBAD ΔaraFGH Φ(ΔaraEp $P_{CP18}$-araE) | Lennen et al. 2010 |
| *E. coli* CM24 | *E. coli* LS5218 ΔfadE ΔatoC ΔldhA ΔackApta ΔadhE ΔpoxB ΔfrdABCD ΔydiO ΔfadBA ΔfadIJ ΔfadD | Mehrer et al. 2018 |

Table 2: Key plasmids

| Name | Description | Source |
|---|---|---|
| pET 28 MA-ACR | ACR from *Marinobacter aqueolei* VT8 on pET 28 backbone, T7 promoter KmR | Rung Yi Lai, (currently Suranaree University of Technology), while a post-doc at UW-Madison |
| pBTRKtrc | Ptrc promoter, pBBR1 origin, KanR | Youngquist et al. 2013 |
| pBTRCK MA-ACR | ACR from *Marinobacter aqueolei* VT8 on pBTRCKtrc backbone | Youngquist et al. 2013 |
| pBTRCK MB-ACR | ACR from *Marinobacter BSs20148* on pBTRCKtrc backbone | Mehrer et al. 2018 |
| pBTRCK MT-ACR | ACR from *Methylibium Sp. T29* on pBTRCKtrc backbone | Mehrer et al. 2018 |
| pACYC-seFadBA | FadBA from *Salmonella enterica*, pACYC origin, Trc promoter, CmR | Mehrer et al. 2018 |
| pTRC99A-vhTER-fdh | Trans-enyol-CoA reductase (TER) from *Vibrio harveyi* and formate dehydrogenase from *Candida boindinii*, pBR322 origin, Trc Promoter, AmpR | Mehrer et al. 2018 |
| pET 28 Ec-ACP | Apo-Acyl Carrier Protein (ACP) from *E. coli* | Hernández-Lozada et al. 2018 |
| pET 28 Vh-AasS | AasS from *V. harveyi* | Hernández-Lozada et al. 2018 |
| pET 28 Bs-Sfp | Sfp from Bascillus subtilis | Hernández-Lozada et al. 2018 |
| pLacIRARE rTEV (pQE60) | TEV Protease | Hernández-Lozada et al. 2018 (originally from Hazel Holden Lab) |

Table 3: Protein structure templates used for homology models

| Template (PDB) | Protein Function | %ID to MA-ACR |
|---|---|---|
| 3m1aA | Short-chain dehydrogenase | 40 |
| 3rkrA | Short-chain oxidoreductase | 37 |
| 3rihA | Putative short-chain dehydrogenase or reductase | 36 |
| 3afmB | Aldose reductase | 36 |
| 3afnB | Aldose reductase | 36 |
| 4bmvA | Short-chain dehydrogenase | 36 |

Table 4: Details of UCB optimization rounds

| | Data used for model training | Model used for design | Sequence design criteria | # sequences designed | # sequences tested | Allowed Block Subs | Classifier Activity Threshold (mg/L) | GNB AUC | Method for Lambda Selection | Lambda | GP Model Correlation Coefficient | Sum of Squared Error |
|---|---|---|---|---|---|---|---|---|---|---|---|---|
| Initialization | None | None | Max MI | 20 | 20 | 5 | N/A | N/A | N/A | N/A | N/A | N/A |
| UCB1 | CM24 + BL21 | GPR with Hamming kernel | UCB | 10 | 5 | 5 | None | None | Inspection | 0.01 | Not Calculated | Not Calculate d |
| UCB2* | RL08ara | NB, GPR with Hamming kernel | UCB positive | 10 | 7 | 5 | 1.1 | Not Calculated | Inspection | 0.433 | 0.192 | 11.46 |
| UCB3 | RL08ara | NB, GPR with Hamming kernel | UCB positive | 10 | 7 | 5 | 1.1 | Not Calculated | Inspection | 0.433 | -0.368 | 38.77 |
| UCB4 | RL08ara | NB, GPR with Hamming kernel | UCB positive | 10 | 5 | 5 | 1.1 | 0.7086 | Maximize Correlation Coefficient | 0.00658 | -0.275 | 23.91 |
| UCB5 | RL08ara | NB, GPR with structure kernel | UCB positive | 10 | 4 | 2 | 1.1 | 0.7472 | Minimize Sum of Squared Error | 0.000167 | 0.189 | 19.2 |
| UCB6** | RL08ara | NB, GPR with structure kernel | UCB positive | 10 | 9 | 2 | 4 | 0.7886 | Minimize Sum of Squared Error | 0.0599 | 0.632 | 5.2 |
| UCB7 | RL08ara | NB, GPR with structure kernel | UCB positive | 10 | 10 | 2 | 4 | 0.81 | Maximize Correlation Coefficient | 0.0359 | 0.384 | 11.17 |
| UCB8 | RL08ara | NB, GPR with structure kernel | UCB positive | 10 | 7 | 2 | 4 | 0.869 | Inspection | 0.0774 | 0.472 | 13.06 |
| UCB9 | RL08ara | NB, GPR with structure kernel | UCB positive | 10 | 10 | 5 | 4 | 0.8799 | Maximize Correlation Coefficient | 0.00774 | 0.466 | 25 |
| UCB10 | RL08ara | NB, GPR with structure kernel | UCB positive | 12 | 9 | 5 | 8 | 0.9757 | Minimize Sum of Squared Error | 0.0215 | 0.786 | 5.47 |

*Sequences in this round were non-UCB-optimal due to a coding error, but the data generated from these sequences was still valuable for training models.

**The labels for two sequences were accidentally swapped during model training

Table 5: Fatty alchohol titers of chimeric ATRs characterized in RL08ara.

| Name | Block Sequence | Average Titer | Number of Replicates |
|---|---|---|---|
| ATR-01 | A-ATAATTBB | 1.5 ± 0.3 | 3 |
| ATR-02 | A-TATTTTAB | 0.8* | 1 |
| ATR-03 | A-TTTTBTBA | 0.9* | 1 |
| ATR-04 | A-ATTBAATB | 0.6* | 1 |
| ATR-05 | A-ABTATTTA | 0.9* | 1 |
| ATR-06 | A-TAATBTTB | 0.9* | 1 |
| ATR-07 | A-AATAATBT | 0.8* | 1 |
| ATR-08 | A-ATTABTAT | 0.8* | 1 |
| ATR-09 | A-TBATATAB | 1.0* | 1 |
| ATR-10 | A-TTBAATTA | 2.8 ± 0.1 | 3 |
| ATR-11 | A-ATBTATTA | 0.7* | 1 |
| ATR-12 | A-BAABTTAA | 0.9* | 1 |
| ATR-13 | A-BTTAAATB | 0.8* | 1 |
| ATR-14 | A-TBBTTABT | 1.0* | 1 |
| ATR-15 | A-TBABATTT | 0.7* | 1 |
| ATR-16 | A-BBATBATT | 0.7* | 1 |
| ATR-17 | A-BBTTATBA | 0.7* | 1 |
| ATR-18 | A-TTABTABA | 2.2 ± 0.2 | 3 |
| ATR-19 | A-TBTABAAA | 0.8* | 1 |
| ATR-20 | A-BTBATTAT | 2.6 ± 0.2 | 3 |
| ATR-21 | A-AABTTAAT | 0.8* | 1 |
| ATR-22 | A-BABBTATA | 1.0* | 1 |
| ATR-23 | A-ATBBBAAT | 3.8 ± 0.3 | 3 |
| ATR-24 | A-TABATABT | 0.8* | 1 |
| ATR-25 | A-BTBTTAAB | 5.0 ± 0.9 | 3 |
| ATR-26 | A-TAAABTBB | 3.6 ± 0.6 | 3 |
| ATR-27 | A-TTBTTTAA | 4.0 ± 1.0 | 3 |
| ATR-28 | A-TBAAATAA | 2.7 ± 0.2 | 3 |
| ATR-29 | A-TBTTATTB | 0.5 ± 0.1 | 3 |
| ATR-30 | A-TAATAATA | 0.6 ± 0.1 | 3 |
| ATR-31 | A-TBTTTATB | 0.6 ± 0.2 | 3 |
| ATR-32 | A-TBTATATA | 0.7 ± 0.2 | 3 |
| ATR-33 | A-BTTAATBA | 1.0 ± 0.2 | 5 |
| ATR-34 | A-BTATBTTA | 5.0 ± 2.0 | 5 |
| ATR-35 | A-ABBATABA | 1.0 ± 0.3 | 5 |
| ATR-36 | A-AABBTTBA | 1.3 ± 0.4 | 5 |
| ATR-37 | A-BABTTTBA | 1.4 ± 0.2 | 5 |

| | | | |
|---|---|---|---|
| ATR-38 | A-AABTTTBA | 1.4 ± 0.3 | 5 |
| ATR-39 | A-AABATABA | 1.1 ± 0.3 | 5 |
| ATR-40 | A-BTBBBABB | 21.0 ± 2.0 | 3 |
| ATR-41 | A-ATAAAAAB | 27.0 ± 7.0 | 3 |
| ATR-42 | A-AABABAAA | 12.0 ± 1.0 | 3 |
| ATR-43 | A-TTBTBTTT | 2.9 ± 0.3 | 3 |
| ATR-44 | A-AAABBAAA | 13.0 ± 1.0 | 2 |
| ATR-45 | A-AAABAAAB | 33.0 ± 6.0 | 3 |
| ATR-46 | A-AABAAAAB | 19.0 ± 2.0 | 3 |
| ATR-47 | A-BBBTBABB | 8.1 ± 0.8 | 3 |
| ATR-48 | A-BAAAAAAB | 16.0 ± 2.0 | 3 |
| ATR-49 | A-ABAAAAAT | 3.6 ± 0.1 | 2 |
| ATR-50 | A-BBBBBATB | 1.4 ± 0.1 | 2 |
| ATR-51 | A-BBBBBTAB | 20.0 ± 3.0 | 2 |
| ATR-52 | A-ABBAAAAA | 12.0 ± 3.0 | 2 |
| ATR-53 | A-ABBBBTBB | 21.0 ± 1.0 | 2 |
| ATR-54 | A-AABAAAAT | 4.4 ± 0.1 | 2 |
| ATR-55 | A-BBBBBTTB | 1.3 ± 0.2 | 2 |
| ATR-56 | A-ATBAAAAA | 14.0 ± 4.0 | 2 |
| ATR-57 | A-BBBABTBB | 9.5 ± 0.6 | 2 |
| ATR-58 | A-AAAABATA | 0.8 ± 0.1 | 3 |
| ATR-59 | A-BBBBBAAB | 49.1 ± 0.9 | 4 |
| ATR-60 | A-ATABAAAA | 19.0 ± 1.0 | 2 |
| ATR-61 | A-AABBAAAA | 12.0 ± 2.0 | 3 |
| ATR-62 | A-BABBBABB | 37.0 ± 5.0 | 2 |
| ATR-63 | A-ABBBBABB | 36.0 ± 4.0 | 3 |
| ATR-64 | A-AAAABAAB | 13.2 ± 0.6 | 2 |
| ATR-65 | A-ABAAAAAB | 14.0 ± 2.0 | 2 |
| ATR-66 | A-BTBBBTBB | 18.8 ± 0.7 | 2 |
| ATR-67 | A-BBBABABB | 7.0 ± 5.0 | 2 |
| ATR-68 | A-BTBBBATB | 7.7 ± 0.4 | 2 |
| ATR-69 | A-BTBBBAAB | 54.0 ± 6.0 | 4 |
| ATR-70 | A-BBBBAAAB | 39.0 ± 4.0 | 2 |
| ATR-71 | A-ABBBBAAB | 45.0 ± 14.0 | 4 |
| ATR-72 | A-BABBBAAB | 34.0 ± 6.0 | 2 |
| ATR-73 | A-BBABBATB | 2.8 ± 0.4 | 2 |
| ATR-74 | A-ATBBBABB | 19.0 ± 2.0 | 2 |
| ATR-75 | A-ABBBBATB | 2.4 ± 0.3 | 2 |
| ATR-76 | A-BABBBATB | 4.3 ± 0.01 | 2 |

| | | | |
|---|---|---|---|
| ATR-77 | A-AABBBABB | 35.5 ± 0.05 | 2 |
| ATR-78 | A-ATBBATAB | 41.0 ± 10.0 | 2 |
| ATR-79 | A-BTABBAAB | 54.0 ± 1.0 | 2 |
| ATR-80 | A-BTBBAAAB | 60.0 ± 4.0 | 3 |
| ATR-81 | A-ATBTBAAB | 24.0 ± 24.0 | 4 |
| ATR-82 | A-BTABAAAB | 42.0 ± 21.0 | 3 |
| ATR-83 | A-ATBBAAAB | 54.0 ± 11.0 | 11 |
| ATR-84 | A-AABBBAAB | 36.0 ± 10.0 | 2 |
| ATR-85 | A-ATABBAAB | 48.0 ± 7.0 | 2 |
| ATR-86 | A-ATBBBAAB | 45.0 ± 19.0 | 3 |
| ATR-87 | A-ATABAAAB | 50.0 ± 12.0 | 2 |
| ATR-88 | A-ATBBBTAB | 33.0 ± 3.0 | 2 |
| ATR-89 | A-AABBAAAB | 40.0 ± 4.0 | 2 |
| ATR-90 | A-BTBAAAAB | 37.0 ± 2.0 | 2 |
| ATR-91 | A-ATBABAAB | 16.0 ± 3.0 | 2 |
| ATR-92 | A-ABBBAAAB | 35.0 ± 4.0 | 2 |
| ATR-93 | A-BTBBATAB | 34.6* | 1 |
| MA-ACR (Parent A) | A-AAAAAAAA | 11.0 ± 3.0 | 26 |
| Parent B (Fusion A-B) | A-BBBBBBBB | 37.0 ± 8.0 | 13 |
| Parent T (Fusion A-T) | A-TTTTTTTT | 6.0 ± 2.0 | 10 |
| MB-ACR | B-BBBBBBBB | 26.0 ± 8.0 | 7 |
| Fusion B-A | B-AAAAAAAA | 11.0 ± 3.0 | 4 |
| Fusion B-T | B-TTTTTTTT | 6.0 ± 2.0 | 4 |
| MT-ACR | T-TTTTTTTT | 4.0 ± 1.0 | 7 |
| Fusion T-A | T-AAAAAAAA | 1.5 ± 0.2 | 4 |
| Fusion T-B | T-BBBBBBBB | 2.1 ± 0.5 | 4 |

*Only single replicate was observed in RL08ara

Titers reported in Table 5 are final titers after all experiments, including additional validation experiments were complete. Because of this, some titers differ from titers shown in the figures of the main text, which reflect titers observed during the UCB optimization process.

Table 6: Amino-acid sequences of blocks and sequence elements

| | Element/ Block | Amino Acid Sequence |
|---|---|---|
| MBP Tag | MBP Tag | MKIEEGKLVIWINGDKGYNGLAEVGKKFEKDTGIKVTVEHPDKLEEKFPQVAATGDGPDIIFWAHDRFGGYAQSGLLAEITPDKAFQDKLYPFTWDAVRYNGKLIAYPIAVEALSLIYNKDLLPNPPKTWEEIPALDKELKAKGKSALMFNLQEPYFTWPLIAADGGYAFKYENGKYDIKDVGVDNAGAKAGLTFLVDLIKNKHMNADTDYSIAEAAFNKGETAMTINGPWAWSNIDTSKVNYGVTVLPTFKGQPSKPFVGVLSAGINAASPNKELAKEFLENYLLTDEGLEAVNKDKPLGAVALKSYEEELVKDPRIAATMENAQKGEIMPNIPQMSAFWYAVRTAVINAASGRQTVDEALKDAQTNSSSNNNNNNNNNNLGIEGRISEF |
| AHR Domains | AHR Domain - A | NYFLTGGTGFIGRFLVEKLLARGGTVYVLVREQSQDKLERLRERWGADDKQVKAVIGDLTSKNLGIDAKTLKSLKGNIDHVFHLAAVYDMGADEEAQAATNIEGTRAAVQAAEAMGAKHFHHVSSIAAAGLFKGIFREDMFEEAEKLDHPYLRTKHESEKVVREECKVPFRIYRPGMVIGHSETGEMDKVDGPYYFFKMIQKIRHALPQWVPTIGIEGGRLNIVPVDFVVDALDHIAHLEGEDGNCFHLVDSDPYKVGEILNIFCEAGHAPRMGMRIDSRMFGFIPPFIRQSIKNLPPVKRITGALLDDMGIPPSVMSFINYPTRFDTRELERVLKGTDIEVPRLPSYAPVIWD |
| | AHR Domain B | NYFVTGGTGFIGRFLIAKLLARGAIVHVLVREQSVQKLADLREKLGADEKQIKAVVGDLTAPSLGLDKKTLKQLSGKIDHFFHLAAIYDMSASEESQQAANIDGTRAAVAAAEALEAGIFHHVSSIAVAGLFKGTFREDMFAEAGKLDHPYFRTKHESERVVRDDCKVPFRIYRPGLVIGDSATGDMDKVDGPYYFFKMIQKIRGALPQWVPTIGIEGGRLNIVPVNFVADALDHIAHLPNEDGKCFHLVDSDPYKVGEILNIFCEAGHAPKMGMRIDSRMFGFVPPFIRQSLKNLPPVKRMGRALLDDLGIPASVLSFINYPTRFDARETERVLQGTGIEVPRLPDYAPVIW |
| | AHR Domain T | QYFVTGATGFIGKRLVRKLLDRRGSTVHFLLRPESERKLPELLAYWGLSGAAKARAVPVYGDLTAKKLGVAADAIKALKGRIDAIYHLAAVYDLGADEAAQVQVNIEGTRSAVEFAQAIQAGHFHHVSSIAAAGLYEGVFREDMFDEAEGLDHPYFMTKHESEKIVRKECKLPWTVFRPAMVVGDSTTGEMDKIDGPYYFFKLIQRMRQLLPPWMPAVGLEGGRVNIVPVDFVVAALDHISHAKLELDRRCFHLVDPVGYRVGDVLDIFGKAAHAPKMNLFVNAALLGFIPKSVKKGLMALAPVRRIRNAVMKDLGLPEDMLTFVNYPTRFDCRDTQAALKGSGIECPNLKDYAWRLW |
| ATR Domain Blocks | 1A | WERNLDPDLFKDRTLKGTVEGKVCV |
| | 1B | WERNLDPDLFKDRTLRGTVEGKVCV |
| | 1T | WERNLDPDLFIDRSLRGTVGGKVVL |
| | 2A | VTGATSGIGLATAEKLAEAGAILVIGARTKETLDEVAASLEAKGGNVHAYQCDFS |
| | 2B | VTGATSGIGLATAEKLADAGAILVIGARTQETLDQVSAQLNARGADVHAYQCDFA |
| | 2T | VTGGSSGIGLAAACKFAEAGAVTVICARDADKLDEAVKEIKAFAGKEARVFSYSVDIA |
| | 3A | DMDDCDRFVKTVLDNHGHVDVLVNN |
| | 3B | DMDACDRFIQTVSENHGAVDVLINN |
| | 3T | DEAGCKAFLEALQAEHGGVDFLINN |
| | 4A | AGRSIRRSLALSFDRFHDFERTMQLNY |
| | 4B | AGRSIRRSLDKSFDRFHDFERTMQLNY |
| | 4T | AGRSIRRAIENSYERFHDFERTMQLNY |
| | 5A | FGSVRLIMGFAPAMLERRRGHVVNISSIGVLTNAPRFSAYVSSKSALDAFSRCAAAEWSDRNVTF |
| | 5B | FGSLRLIMGFAPAMLERRRGHIINISSIGVLTNSPRFSAYVASKSALDSFSRCAAAEWSDRRVCF |
| | 5T | FGCLRVTMGVLPGMVAKRKGHVVNISSIGVLTNAPRFSAYVASKAALDAWTRCASSEYADTGISF |
| | 6A | TTINMPLVKTPMIAPTKIYDSVPT |

| | | |
|---|---|---|
| | **6B** | TTINMPLVKTPMIAPTKIYDSVPT |
| | **6T** | TTINMPLVRTPMIAPTKIYNNVPT |
| | **7A** | LTPDEAAQMVADAIVYRPKRIATRLGVFAQVLHALAPKMGEIIMNTGYRM |
| | **7B** | LSPEEAADMVVNAIVYRPKRIATRMGVFAQVLNAVAPKASEILMNTGYKM |
| | **7T** | LAPEEAADMIAQACVYKPVRIATRLGTAGQVLHALAPRVAQIVMNTSFRM |
| | **8A** | FPDSPAAAGSKSGEKPKVSTEQVAFAAIMRGIYW* |
| | **8B** | FPDSMPKKGKEVSAEKGASTDQVAFAAIMRGIHW* |
| | **8T** | FPDSEAAKGEKGAKPQLSAEAVALQQMMRGIHF* |

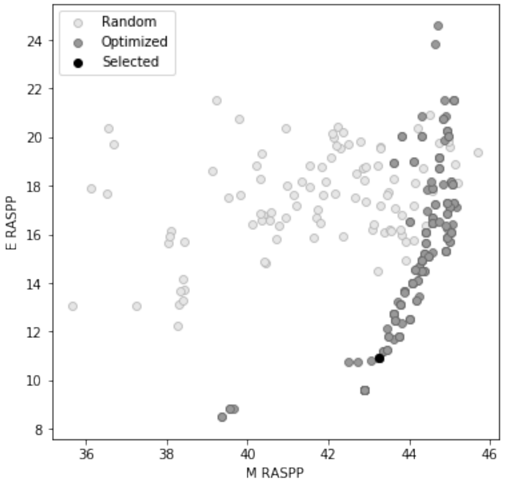

**Figure S1**: RASPP chimeric enzyme library design. The SCHEMA-RASPP algorithm was used to identify sets of breakpoints that simultaneously maximize the average mutation level (M) of the library and minimize the SCHEMA energy (E) of the library. Each point in the graph represents a library of chimeras, and the library that was selected is shown in black. Libraries with randomized breakpoints are shown in light gray for comparison.

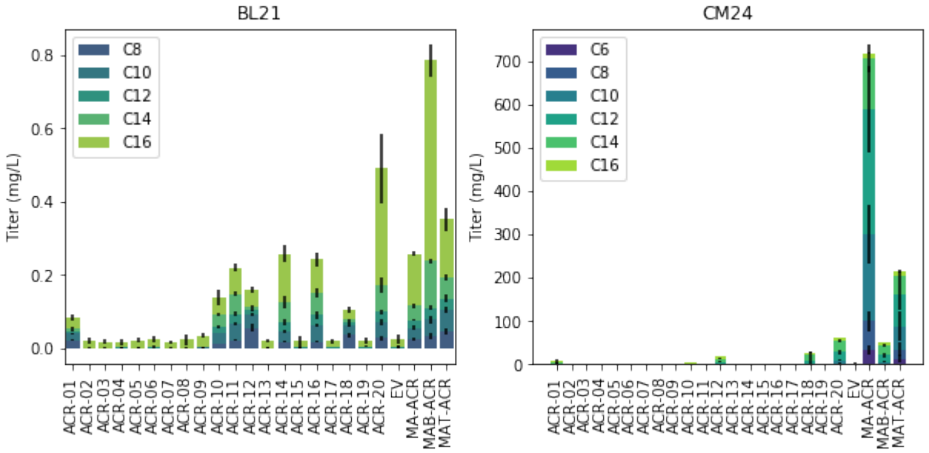

**Figure S2**: Fatty alcohol titer data in BL21 (DE3) and CM24 for the initial chimera seed sample. The total titers from these experiments were used to train the models used for the first round of UCB optimization.

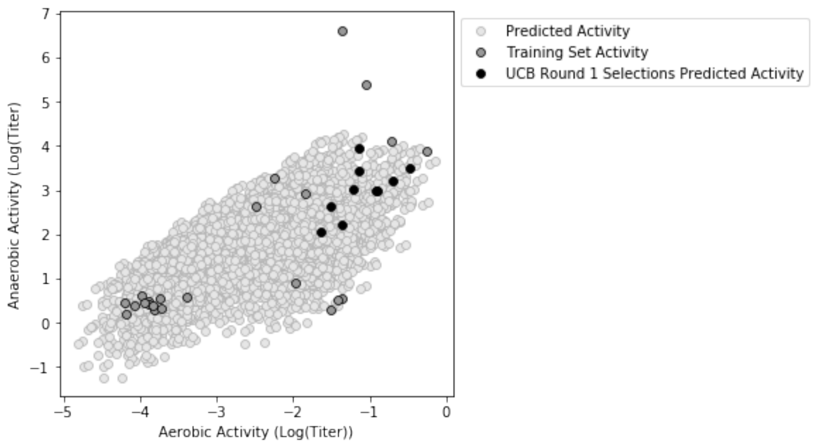

**Figure S3**: Model predictions for UCB1. Models trained on BL21 data (aerobic) and CM24 data (anaerobic) were compared and used to determine a set of sequences that optimized both UCBs.

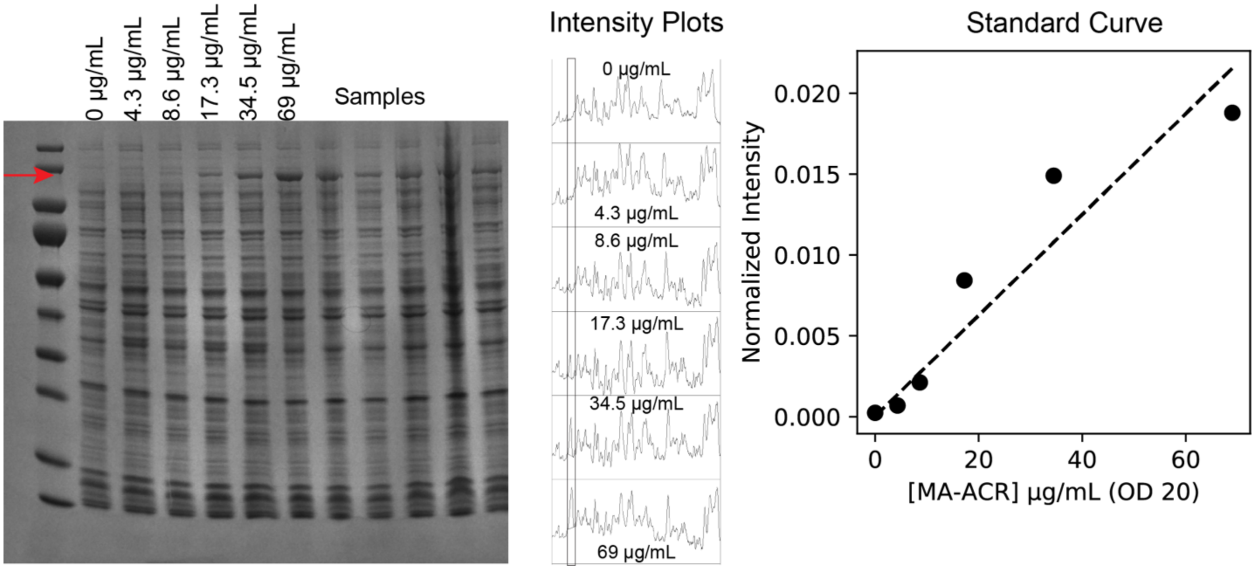

**Figure S4**: Representative SDS-PAGE gel for measuring FAR expression level, image analysis, and standard curve.

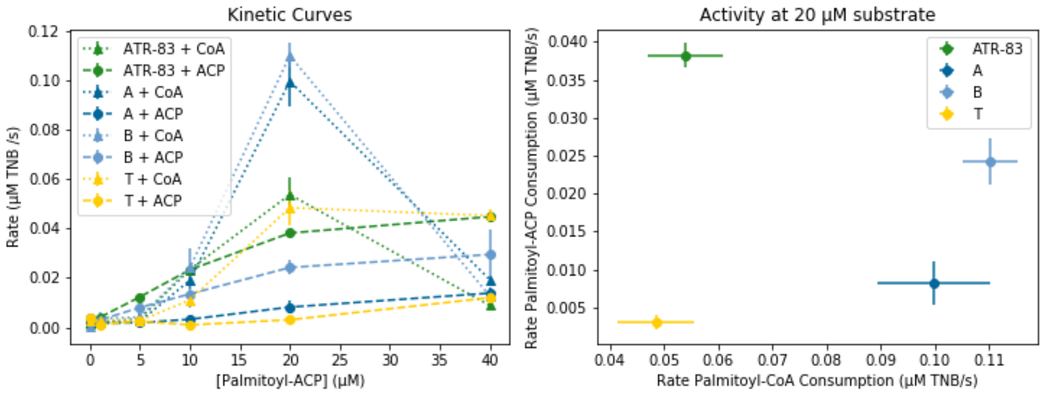

**Figure S5**: Comparison of enzyme activity on palmitoyl-CoA substrates

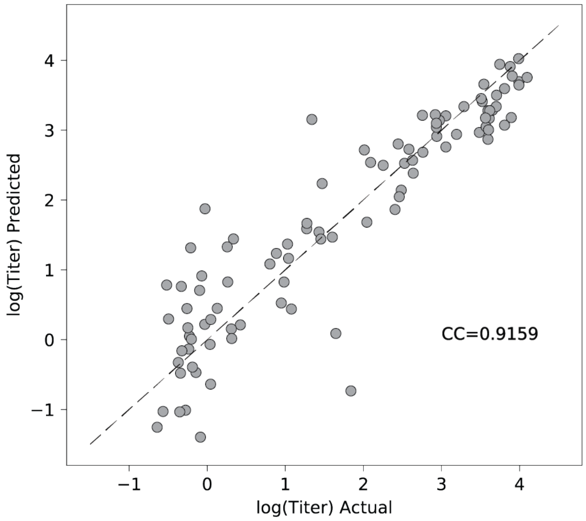

**Figure S6**: Cross-validated Gaussian process regression model used to study chimera landscape and determine contributions of blocks.
